# Cortical VIP neurons locally control the gain but globally control the coherence of gamma band rhythms

**DOI:** 10.1101/2021.05.20.444979

**Authors:** Julia Veit, Gregory Handy, Daniel P. Mossing, Brent Doiron, Hillel Adesnik

**Author notes:** Department of Physiology, University of Freiburg, Freiburg, Germany (Julia Veit).

## Abstract

Gamma band synchronization can facilitate local and long-range communication in neural circuits. In the primary visual cortex (V1) the strength of synchronization on the local level is strongly tuned to the contrast, size and center/surround orientation of grating stimuli. On the global level, the synchronization of gamma oscillations across the retinotopic map crucially depends on matched stimulus properties in the corresponding locations in the visual field. Although these features of V1 gamma rhythms are likely to be crucial for how they might support cortico-cortical communication and visual perception, their neural basis remains largely unknown. We hypothesized VIP disinhibitory interneurons, which shape other tuning properties in V1 by inhibiting SST neurons, may be responsible for tuning local gamma band power and global gamma synchronization. To test these ideas, we combined multi-electrode electrophysiology, cell-type specific optogenetic suppression of VIP neurons and computational modeling. Contrary to expectations, our data show that on the local level, VIP activity has no role in tuning gamma power to stimulus properties; rather, it scales the gain of gamma oscillations linearly across stimulus space and across behavioral state. Conversely, on the global level, VIP neurons specifically suppress gamma synchronization (as measured by spectral coherence) between spatially separated cortical ensembles when they are processing non-matched stimulus features. A straightforward computational model of V1 shows that like-to-like connectivity across retinotopic space, and specific, but powerful VIP➔SST inhibition are sufficient to capture these seemingly opposed effects. These data demonstrate how VIP neurons differentially impact local and global properties of gamma rhythms depending on the global statistics of the retinal image. VIP neurons may thus construct temporal filters in the gamma band for spatially continuous image features, such as contours, to facilitate the downstream generation of coherent visual percepts.

## Introduction

Synchronized activity is widespread in neural systems, occurring both spontaneously and during sensory stimulation, cognition, and motor action (Jasper & Penfield, 1949; Adrian, 1950; Bressler & Freeman, 1980; Riehle *et al.*, 1997; Buzsáki & Draguhn, 2004; Colgin *et al.*, 2009; Fries, 2009; Vinck *et al.*, 2015). In monkeys, synchronization is dependent on stimulus features (Gieselmann & Thiele, 2008; Ray & Maunsell, 2010; Ray *et al.*, 2013) and modulated by behavioral state, such as directed attention (Fries *et al.*, 2001; Womelsdorf & Fries, 2006; Chalk *et al.*, 2010; Vinck *et al.*, 2013). Synchronization may facilitate neural communication by enhancing the temporal co-incidence of synaptic excitatory potentials in target neurons (Colgin *et al.*, 2009; Fries, 2015). Gamma band synchrony across distant sites in the primary visual cortex (V1) depends on matched stimulus properties processed by the two sites (Gray & Singer, 1989; Fries, 2009), suggesting a role in promoting the contextual synthesis of visual percepts downstream. Thus, the tuning of local gamma band power and global gamma band coherence to specific stimuli and contexts may be crucial for its role in cortical computation and perception. Imporantly, not only must gamma rhythms be tuned, they must also be scaled appropriately, as excessive synchrony can limit information carrying capacity of neural networks (Benda *et al.*, 2006; Nandy *et al.*, 2019), and too much or too little synchrony may lead to neurological disorders (Lewis *et al.*, 2005; Schnitzler & Gross, 2005; Uhlhaas & Singer, 2010; Yizhar *et al.*, 2011). Remarkably, despite detailed knowledge of the phenomenology of cortical oscillations on one hand, and a deep mechanistic and theoretical insight into their underlying synaptic basis on the other (Traub *et al.*, 2004; Bartos *et al.*, 2007; Buzsáki & Wang, 2012) we have a very limited understanding for the neural circuits that regulate their magnitude and coherence across different sensory and behavioral contexts (Fries *et al.*, 2001; Chalk *et al.*, 2010).

Mechanistically, ample evidence indicates that local GABAergic interneurons temporally entrain excitatory neurons by biasing their spike timing to the trough of their periodic inhibitory synaptic potentials (Bartos *et al.*, 2002, 2007; Hasenstaub *et al.*, 2005, 2016; Tukker *et al.*, 2007; Wulff *et al.*, 2009; Buzsáki & Wang, 2012; Perrenoud *et al.*, 2016; Zhang *et al.*, 2018). This periodicity results from the recurrent interaction between excitatory and inhibitory neurons (Hasenstaub *et al.*, 2005), through direct interneuron-to-interneuron synaptic coupling (Sohal & Huguenard, 2005), and through electrical synapses (Traub *et al.*, 2001; Long *et al.*, 2005; Neske & Connors, 2016). Cortical gamma oscillations depend on various types of interneurons, including soma-targeting parvalbumin positive basket cells (Cardin *et al.*, 2009; Sohal *et al.*, 2009). In the mouse primary visual cortex, a visually induced gamma oscillation (25-40 Hz), similar to the widely studied gamma rhythms in higher mammals (Gray *et al.*, 1989; Gieselmann & Thiele, 2008; Ray & Maunsell, 2010; Self *et al.*, 2016), requires the activity of somatostatin (SST) interneurons (Chen *et al.*, 2017; Veit *et al.*, 2017; Hakim *et al.*, 2018). In V1, SST neuron firing rates strongly correlate with visually induced narrow-band gamma power on a trial-to-trial basis, and optogenetic inactivation of SST neurons (but not PV neurons) nearly abolishes visually evoked gamma oscillations (Veit *et al.*, 2017). SST neurons are also known to be critical for the encoding of contextual stimuli, such as for gratings that extend beyond neurons’ classical receptive fields (Adesnik *et al.*, 2012; Nienborg *et al.*, 2013; Keller *et al.*, 2020; Mossing *et al.*, 2021). Notably, a second narrowband gamma oscillation around ~60 Hz that is increased by locomotion and luminance, but strongly suppressed by visual stimuli, is also present in V1 but is not of cortical origin and thus independent of cortical interneurons (Saleem *et al.*, 2017; Storchi *et al.*, 2017; Veit *et al.*, 2017; Hoseini *et al.*, 2021).

The discovery that VIP interneurons preferentially inhibit other interneurons, especially SST neurons, (Pfeffer *et al.*, 2013; Pi *et al.*, 2013; Karnani, Jackson, Ayzenshtat, Hamzehei Sichani, *et al.*, 2016) raises the hypothesis that they might regulate the power and stimulus-dependence of gamma band oscillations. Recent work has shown that VIP neurons are suppressed by visual stimuli (Keller *et al.*, 2020; Millman *et al.*, 2020; Mossing *et al.*, 2021) that have previously been shown to drive strong gamma oscillations (Gieselmann & Thiele, 2008; Ray *et al.*, 2013; Veit *et al.*, 2017), but no direct link has been established. In particular, large, high contrast and iso-oriented gratings potently drive gamma rhythms in V1, but simultaneously supress VIP activity. Conversely, small, low contrast, cross-oriented gratings weakly induce gamma rhythyms, but drive strong VIP firing. These results raise the hypothesis that while SST neurons induce gamma rhythms by inhibiting pyramidal cells, VIP neurons tune the gamma rhythm to specific stimulus features by modulating SST neurons. If so, this would establish VIP-mediated disinhibition as a crucial regulator of local gamma band synchronization.

In this study we test whether VIP neurons play a role in tuning gamma power and synchrony globally across the retinotopic map (hereafter termed global coherence) via their disinhibitory action in the cortical microcircuit. Specifically, if VIP neurons suppress SST neurons they would reduce local gamma power and this may also serve to reduce global coherence. However, the efficacy with which VIP neuron activity suppresses global coherence may depend upon whether the stimulus features in distant sites conflict or match with one another. To test this idea we combined multi-site, multi electrode array recordings and optogenetics in awake mice with computational modeling of the superficial layers of the V1 network. We found that VIP suppression only scaled the gain of local gamma band synchronization, but did not alter its tuning, contrary to expectations. Conversely, we found that VIP suppression enhanced the global coherence between distant sites in V1 preferentially when those sites were processing non-matched stimulus features. Remarkably, VIP activity could simultaneously suppress gamma power locally but selectively permit gamma coherence globally for large, homogenous textures. This demonstrates a stimulus-dependent decoupling between the local and global properties of gamma oscillations. A computational model of L2/3 in mouse V1 captures all of these findings with only the minimal conditions of like-to-like connectivity across space and selective inhibition of SST by VIP. The ubitiquity of these features throughout cortex suggest that our findings may generalize beyond the visual system.

These results reveal contrasting local and global circuit roles of VIP-mediated disinhibition in the visual cortex. They demonstrate that locally, VIP activity regulates the gain, but not the tuning of gamma power, while globally VIP neurons contribute to feature-dependent network synchronziation. These widespread effects of VIP neuron suppression might help explain why perturbation of VIP neurons – whether it be genetically, pharmacologically, or optogenetically – potently impairs visual behaviors and learning. Furthermore, they raise the notion that developmental defects in VIP neurons might lead to a range of neurological disorders that have been linked to changes in cortical rhythms, potentially through maladaptive hyper-synchronization.

## Results

### VIP neurons locally control the gain but not the tuning of visually induced gamma oscillations in V1

In primates, cats, and humans, visual stimuli can induce potent oscillations in the gamma band (20-90 Hz), yet the strength of these rhythms depends on the properties of the visual stimulus (Gray *et al.*, 1989; Gieselmann & Thiele, 2008; Ray & Maunsell, 2010; Hermes *et al.*, 2015; Self *et al.*, 2016; Bartoli *et al.*, 2019; Peter *et al.*, 2019) and the brain state (Chalk *et al.*, 2010; Bosman *et al.*, 2012). To probe the neural mechanisms of gamma rhythms in mouse primary visual cortex (V1), we presented head-fixed, awake, locomoting mice with drifting gratings varying in size, contrast, or the orientation of the grating surround relative to the center. We inserted one or two laminar multielectrode arrays into the superficial layers of the primary visual cortex to record both isolated single units and the local field potential (LFP) and used optogenetic perturbations to probe the underlying circuit mechanisms (Fig. 1A). Visual stimuli potently and specifically induced narrow band gamma oscillations (~30 Hz), and gamma power rose monotonically with stimulus contrast and size, (Fig 1 B,C; Fig. S1 A-C), but decreased as the relative angle of orientation between the center and surround was increased (Fig 1 D; Fig. S1 D). These effects in the LFP were mirrored by those in the phase-locking of isolated single units (Fig. S2) confirming that they were mediated by local changes in spike synchrony. Gamma power across these stimulus dimensions was also strongly modulated by behavioral state (Fig. S3).

**Figure 1:**
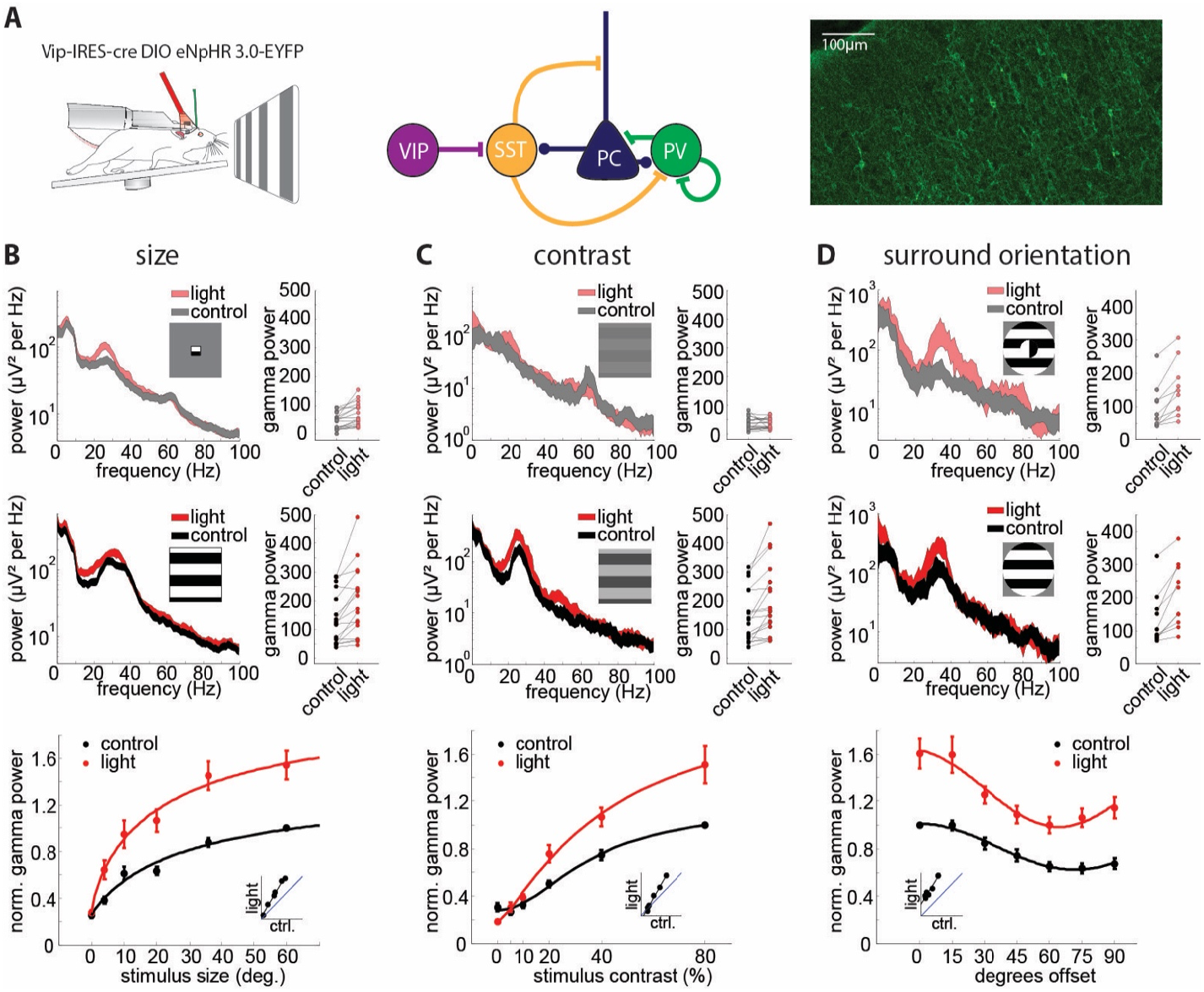
VIP neurons locally control the gain but not the tuning of visually induced gamma oscillations. **A:** Left: Schematic of a head-fixed mouse on a running wheel with an optic fiber over the visual cortex and a laminar multi-electrode array in V1. Middle: Simplified circuit diagram with VIP neurons disinhibiting PCs from SST inhibition. Right: Example image of a V1 brain section from a VIP-Cre mouse injected with a Cre-dependent AAV virus driving eNpHR3.0-YFP. **B:** Top: Left: example LFP power spectrum in response to a small (4°) drifting grating with (red hue) and without (gray) light mediated suppression of VIP neurons (thickness of line denotes mean ± standard error). Right: plot comparing the gamma power to small gratings with and without light (n = 17, p = 0.0003, Wilcoxon signed rank test). Middle: Left: example LFP power spectrum in response to a large (60°) drifting grating with (red) and without (black) light mediated suppression of VIP neurons (thickness of line denotes mean ± standard error). Right: plot comparing the gamma power to large gratings with and without light (n = 17, p = 0.0008, Wilcoxon signed rank test). Bottom: Average normalized gamma power with (red) and without (black) optogenetic suppression of VIP neurons versus stimulus size (n = 17, 2-way-ANOVA: main effect of light: F(1,160) = 54.18, p<0.001; main effect of size: F(4,160) = 22.18, p<0.001; interaction: F(4,160) = 1.03, p = 0.39). **C:** Top: Left: example LFP power spectrum in response to a low contrast (5%) drifting grating with (red hue) and without (gray) light-mediated suppression of VIP neurons (thickness of line denotes mean ± standard error). Right: plot comparing the gamma power to low contrast gratings with and without light (n = 18, p = 0.45, Wilcoxon signed rank test). Middle: Left: example LFP power spectrum in response to a high contrast (80%) drifting grating with (red) and without (black) light-mediated suppression of VIP neurons (thickness of line denotes mean ± standard error). Right: plot comparing the gamma power to high contrast gratings with and without light (n = 18, p = 0.0005, Wilcoxon signed rank test). Bottom: average normalized gamma power with (red) and without (black) optogenetic suppression of VIP neurons versus stimulus contrast (n = 18, 2-way-ANOVA: main effect of light: F(1,170) = 27.81, p<0.001; main effect of contrast: F(4,170) = 65.08, p<0.001; interaction: F(4,170) = 3.85, p = 0.005).**D:** Top: Left: example LFP power spectrum in response to a cross surround (90° offset) drifting grating with (red hue) and without (gray) light mediated suppression of VIP neurons (thickness of line denotes mean ± standard error). Right: plot comparing the gamma power to cross surround gratings with and without light (n = 10, p = 0.002, Wilcoxon signed rank test). Middle: Left: example LFP power spectrum in response to an iso surround (0° offset) drifting grating with (red) and without (black) light mediated suppression of VIP neurons (thickness of line denotes mean ± standard error). Right: plot comparing the gamma power to iso surround gratings with and without light (n = 10, p = 0.002, Wilcoxon signed rank test). Bottom: Average normalized gamma power with (red) and without (black) optogenetic suppression of VIP neurons versus relative surround orientation (n = 10, 2-way-ANOVA: main effect of light: F(1,125) = 119.37, p<0.001; main effect of orientation: F(6,125) = 13.14, p<0.001; interaction: F(6,125) = 0.88, p = 0.51).

For three reasons, we hypothesized that cortical VIP neurons might be crucial for the strong feature and behavioral dependence of visually induced gamma rhythms. First, VIP neurons potently inhibit SST neurons (Pfeffer *et al.*, 2013) and SST activity has been shown to be critical for visually driven gamma oscillations in V1 (Veit *et al.*, 2017). Second, VIP neurons control cortical gain across behavioral states (Lee *et al.*, 2013; Fu *et al.*, 2014; Jackson *et al.*, 2016) similar to what we observed for gamma power. Third, across visual stimuli, VIP neurons’ activity has been reported to be lowest when we find gamma power to be the highest: VIP neurons are suppressed by high contrast, suppressed by large gratings, and suppressed by iso-oriented as compared to cross-oriented gratings (Keller *et al.*, 2020; Millman *et al.*, 2020; Mossing *et al.*, 2021). To probe this last notion directly, we correlated gamma power measured electrophysiologically to average SST and VIP neuron activity measured with two-photon imaging in a separate set of mice. While SST neuron activity was highest in conditions that showed high gamma power (R: 0.76, p: 0.019), VIP neuron activity was lowest (R: −0.84, p: 0.005) and vice versa (Fig. S4).

These results raise the hypothesis that VIP neurons might actively tune gamma power to the contrast, size and surround orientation of gratings. If so, this would be in line with recent reports that argue that VIP neurons actively tune the firing rates of pyramidal cells across contrast (Millman *et al.*, 2020) and center/surround orientation (Keller *et al.*, 2020). To test this notion, we optogenetically suppressed VIP neurons via Cre-dependent expression of the potent optogenetic silencer eNpHR3.0. Post-mortem histological analysis revealed widespread expression of eNpHR3.0 in superficial interneurons with bipolar morphology (Figure 1A, right). Illumination of the visual cortex in these mice resulted in significant enhancements in narrowband gamma power (20-40 Hz) across visual stimulus size, contrast, and center/surround orientation (Figure 1B-D). Strikingly, VIP neuron suppression multiplicatively enhanced gamma power across all feature dimensions, thereby scaling the gain of neural synchronization while preserving the tuning to contrast, size and center/surround orientation dependence (Fig. 1B-D bottom). Thus, contrary to expectations based on these prior studies, our results negate the hypothesis that VIP activity generates the strong feature-dependence of gamma band synchronization in V1. These effects in the LFP were mirrored by changes in the phase locking of isolated single unit spiking activity (Fig. S5,6).

Optogenetically suppressing VIP neurons had no effect on the higher frequency (55-65 Hz) narrowband gamma oscillation derived from sub-cortical circuits, demonstrating a specific role of VIP in controlling stimulus-induced cortical gamma synchronization only (Fig. S7).

Since locomotion also controls the gain of visually evoked activity in V1 (Niell & Stryker, 2010) and potently regulates visually induced gamma oscillations (Fig. S3), we asked whether VIP neurons contribute to the behavioral dependence of gamma band synchronization. We found that suppressing VIP neurons strongly enhanced gamma band power and phase coupling of V1 units across both locomoting and quiescent states (Fig. S8A-C, 3-way ANOVA with factors light, locomotion and stimulus with post-hoc testing, all p<0.001, no interactions), but preferentially during locomotion (Fig. S8D-F), demonstrating that VIP neurons regulate the behavioral modulation of network synchronization in mouse V1.

Taken together, these results demonstrate that locally, VIP neurons scale the power of gamma band rhythms both as a function of the stimulus and as a function of brain state. Yet contrary to expectations, they appear to play no role in the tuning of gamma band power to any of these features.

### VIP neurons globally tune the coherence of V1 ensembles

One of the most striking features of V1 gamma band oscillations is that they preferentially synchronize distant neurons that are processing separate parts of a stimulus with common properties, such as the same orientation and direction of motion (Gray *et al.*, 1989), indicative of belonging to a common object. The mechanisms for this fundamental phenomenon remain largely unkown. Optogenetically suppressing SST neurons strongly reduces this global coherence (Veit *et al.*, 2017), but this could be a simple consequence of the strong supporession of local gamma rhyhtms at both locations. One possibility is that VIP neurons preferentially suppress global coherence when the stimulus features between two retinotopic locations conflict, but permit coherence when those features are shared. To test this, we placed one multielectrode array in the retinotopic region corresponding to the center of the grating, and one in a distal region representing the surround (Fig. 2A, average electrode separation: 530+-90 μm n = 7, 15±3 visual degrees n = 11, Fig. S9). Similar to findings in cats (Gray *et al.*, 1989), large, homogeneous (‘iso-oriented’) drifting gratings drove highly coherent LFP gamma oscillations between the two separate sites (Fig. 2B top). However, when the grating orientation for the two separated electrode arrays was orthogonal (‘cross-oriented’), coherence in the gamma frequency band, dropped substantially (Fig. 2B bottom: 23+-5.7%, p = 0.004 Wilcoxon signed rank). If VIP neurons would simply reduce coherence according to the reduction in gamma power, we would expect coherence to increase similarly for both matched (iso) and non-matched (cross) stimulus features. However, optogenetically suppressing VIP neurons had no impact on global coherence for iso-oriented gratings (Fig. 2C,D, 4.5+-2.6%, p = 0.16, Wilcoxon signed rank test), but significantly increased coherence for cross-oriented gratings (18.6+-8.2%, p = 0.008, Wilcoxon signed rank test). The impact on coherence for cross-oriented gratings was highly specific to the visually induced gamma band (~30 Hz) (Fig. 2E,F).

**Figure 2:**
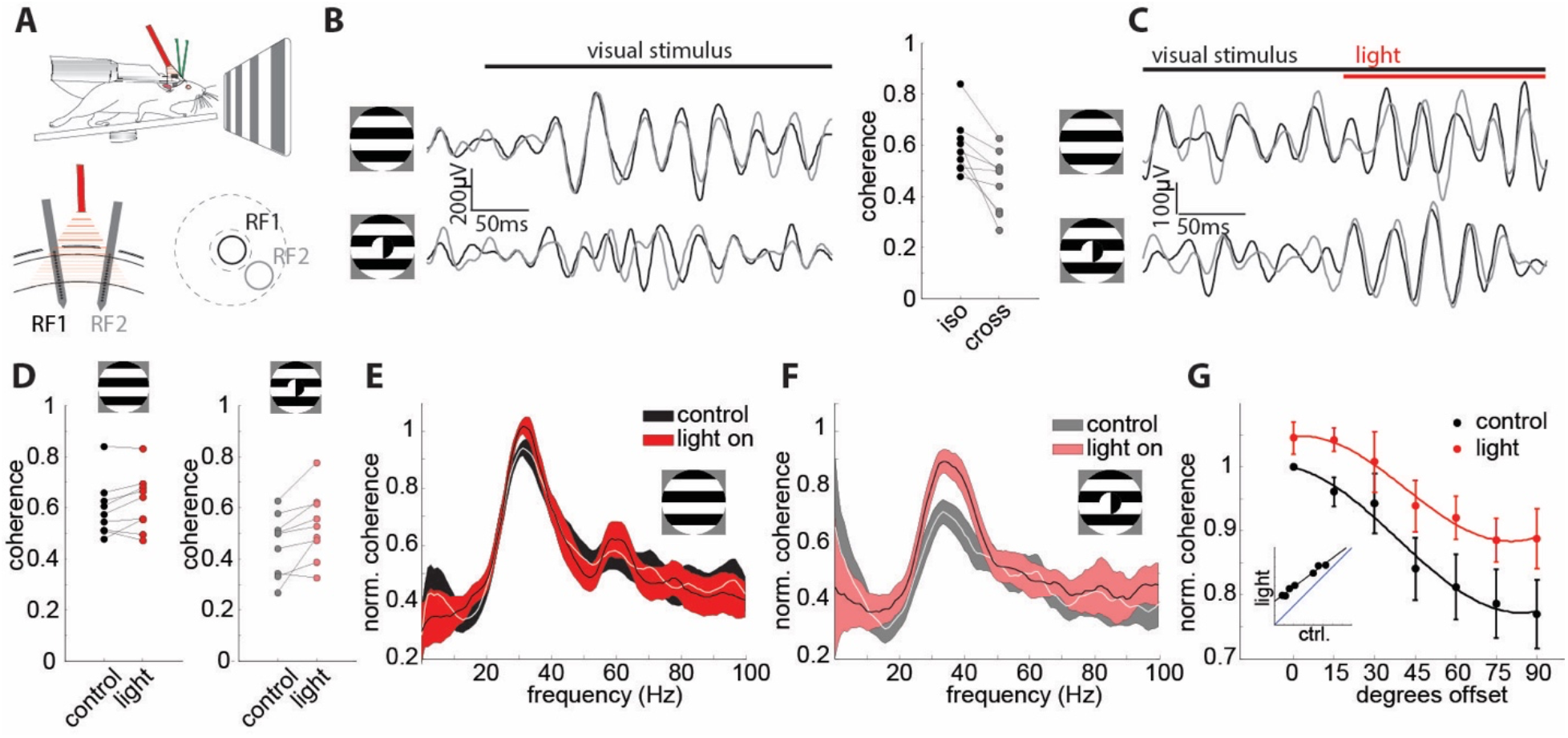
VIP neurons globally tune the coherence of visual ensembles. **A:** Top: recording schematic with two independent laminar probes in V1 of awake, head-fixed VIP-Cre mice. Bottom left: Schematic of the multielectrode array recording configuration with two laminar arrays in distant sites (530+-90 μm apart, histology from n = 7 mice) recorded from two separate receptive fields (RF1 and RF2, 15° ± 3° of visual angle separation, n = 11 mice). Bottom right: schematic of the receptive fields’ locations on the two laminar probes. The center and surround of the gratings are indicated with dashed lines. **B:** Left: Example filtered LFP traces in response to an iso (0° offset, top) and a cross (90° offset, bottom) oriented surround relative to the center. Traces from the center recording site are plotted in black, traces from the surround in gray. Right: Plot comparing the LFP gamma band coherence for iso-oriented to cross-oriented surround stimuli (n = 9, p = 0.004, Wilcoxon signed rank test). **C:** Example filtered LFP traces in response to an iso (0° offset, top) and a cross (90° offset, bottom) oriented surround relative to the center. Traces from the center recording site are plotted in black, traces from the surround in gray. The onset of light to suppress VIP cell activity is shown as a red bar on top. **D:** Left: Plot comparing the LFP gamma band coherence for iso-oriented surround stimuli for control (black) and VIP inactivation (red) trials (n = 9, p = 0.16, Wilcoxon signed rank test) Right: Plot comparing the LFP gamma band coherence for cross-oriented surround stimuli for control (gray) and light (light red) trials (n = 9, p = 0.008, Wilcoxon signed rank test). **E:** Population averaged coherence spectra, normalized to the maximum of control condition, for iso-oriented surround stimuli for control (black) and light (red) trials (n = 10, thickness of line denotes mean ± standard error). **F:** Population averaged coherence spectra for cross-surround stimuli for control (gray) and light (light red) trilas (n = 10, thickness of line denotes mean ± standard error). **G:** Plot of average normalized coherence versus relative surround orientation with (red) and without (black) inactivation of VIP neurons (n = 9, 2-way ANOVA: main effect of light: F(1,107) = 18.8, p<0.001; main effect of offset angle: F(6,107) = 9.16, p<0.001; interaction: F(6,107) = 0.22, p = 0.97). Error bars represent s.e.m.

These results demonstrate that VIP neurons critically contribute to long-range synchronization of primary visual cortical ensembles: they preferentially suppress synchrony when the stimulus features for distant ensembles do not match. Remarkably, even though VIP neuron suppression profoundly enhanced local gamma power in response to iso-oriented gratings (see Fig. 1D), it did not significantly increase coherence. Note that the lack of an increase in coherence for iso-oriented stimuli was not due to a ceiling effect, as the measured coherence was substantially less than the theoretical maximum of 1 (mean coherence iso: 0.60+/−0.04, mean coherence iso+light: 0.62+/−0.04 iso)(Fig. 2C). This demonstrates that the local and global properties of stimulus induced gamma oscillations in the visual cortex can be uncoupled: VIP neurons generally suppress the strength of gamma rhythms for all stimuli, but they only suppress the coherence of gamma rhythms when the features being processing by two distant sites in V1 conflict.

### Computational modeling of V1 explains VIP neurons role in local and global gamma synchronization

To gain insight into how VIP neurons might scale gamma power locally but regulate gamma coherence globally we developed a computational model of layer 2/3 of mouse V1 composed of its four primary cell types modeled by integrate-and-fire spiking dynamics (Fig. 3A). Neurons were connected according to well-known rules (see Supplemental Methods), though to maintain simplicity, VIP neurons only targeted SST neurons and were themselves driven exclusively by an external (untuned) bias input. To account for stimulus size, multiple discrete retinotopic circuits were connected via horizontal excitatory connections, the strength of which was larger for iso-tuned center and surround populations compared to cross-tuned populations (Fig. 3A). The model produced stochastic spiking dynamics that readily generated robust population gamma rhythms (Fig. 3B). The summed population activity was well-captured by a mean-field approximation which is constructed from linearized neuronal responses (Fig. 3C-D, compare the simulation points to the smooth lines computed from the theory; see Supplemental Methods). Despite only including a subset of key features of the mouse V1 network, these gamma rhythms scaled with stimulus strength (≈contrast), size, and center/surround orientation in a qualitatively similar fashion as our experimental results (Fig. 3E-G, gray curves). Moreover, reducing VIP activity to mimic the effects of optogenetically suppressing VIP neurons yielded qualitatively similar impacts on gamma power across stimulus dimensions (Fig. 3E-G, red curves). This shows that a fairly minimal model captures the core phenomenology of visually induced gamma rhythms in V1 and could qualitatively predict the scaling of gamma power across stimulus space by VIP neurons.

**Figure 3:**
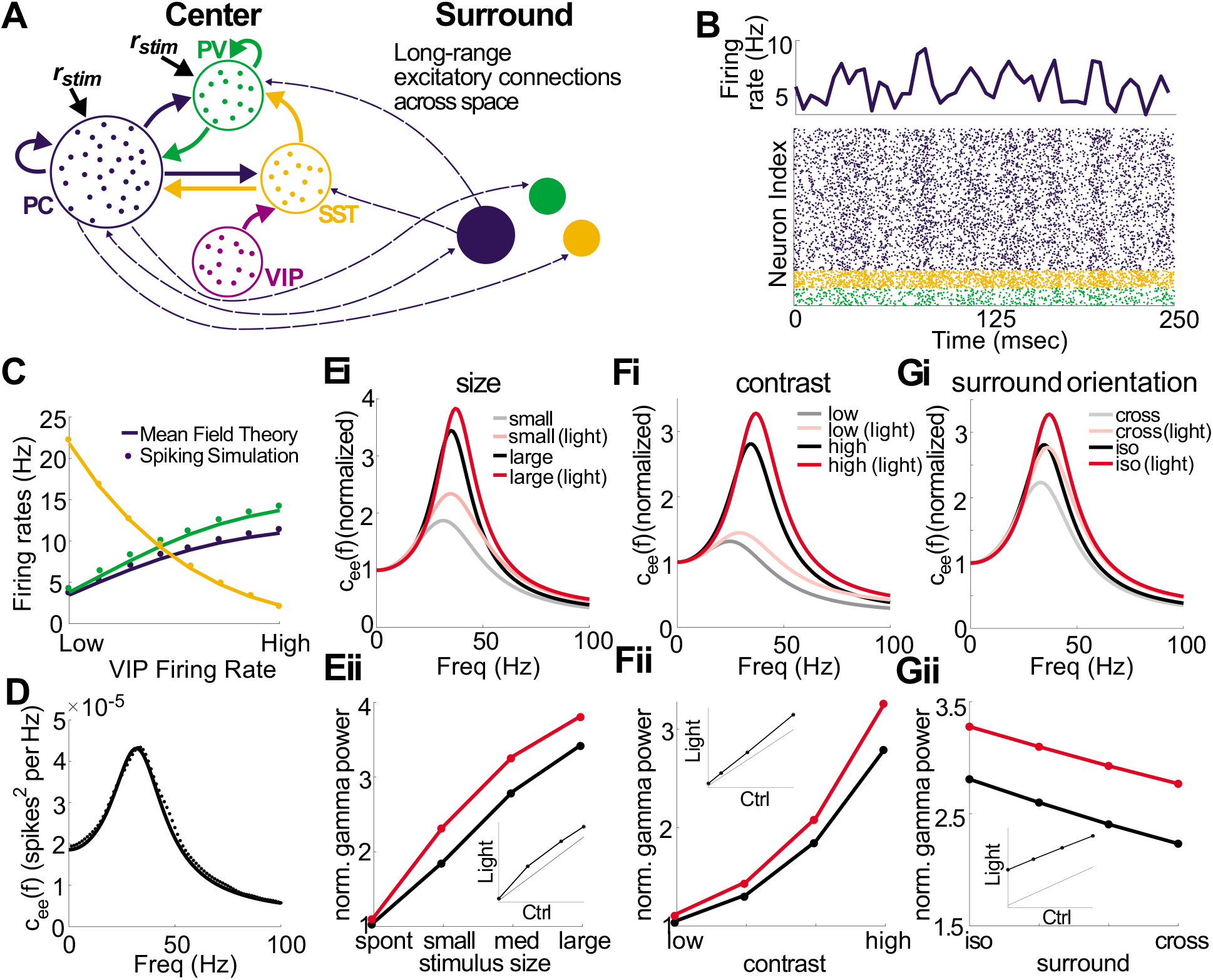
Minimal computational model captures VIP neurons’ role in controlling gamma power. **A:** Schematic of the local connectivity across the four cell types, along with the long-ranged excitatory connections (dashed arrows) spanning across space. All populations receive a static background current, while PC and PV neurons receive stimulus-dependent drives. For a large stimulus size, an additional surround population was added (see Supplemental Methods). **B:** Bottom: Raster plot showing the spike times of neurons from the PC, PV, and SST populations. Top: Average firing rate (averaged over a 5 msec time window) across the excitatory population. **C:** Average firing rates across the populations for the spiking simulation and mean-field theory as a function of VIP firing rates. D: Example of the power spectrum for spiking simulations (dots) and mean-field theory (solid line) showing a strong peak in the gamma frequency. **E:** Ei:Normalized power spectrum from the mean-field model as a function of stimulus size. The red lines illustrate the result of suppressing VIP neurons (i.e., mimicking the optogenetic suppression done experimentally). Eii: Normalized gamma power taken from the power spectrum of panel Ei. **F,G:** Same as E, except for contrast and surround orientation. Similar to experimental results, increasing the size (and contrast) of the stimulus results in a noticeable increase in gamma power. Likewise, iso-surround exhibits larger gamma power than cross-surround. Furter, suppressing VIP leads to a linear increase in gamma power across conditions.

Next, we asked in the model how VIP neurons could globally tune coherence despite locally only scaling gamma power. In agreement with our experimental results, we found that high VIP activity in the model specifically suppressed coherence for cross-oriented as compared to iso-oriented gratings (Fig. 4A-B). We then probed which features of VIP connectivity in the model might be important for this result. First, we constructed a model where VIP directly inhibited PV neurons rather than the SST neurons (Fig. 4C). This model could not reproduce the core experimental results, putatively because PV neurons are key stabilizers in this circuit (Bos *et al.*, 2020). As a result, their suppression led to a large increase in excitatory firing rates, resulting in an increase in gamma power and gamma coherence under the iso-oriented condition (Fig. 4Cii). Finally, we constructed a model where VIP neurons non-specifically targeted all other cell types in the circuit. While this model could capture the impact of VIP activity on overall coherence, it could not recapitulate the selective effect on cross-oriented gratings (Fig. 4D). These modeling experiments imply that the selective inhibitory-inhibitory wiring between VIPs and SSTs is central to the feature-dependence of gamma band coherence across V1 as simpler circuits could not robustly reproduce this core phenomenology.

**Figure 4:**
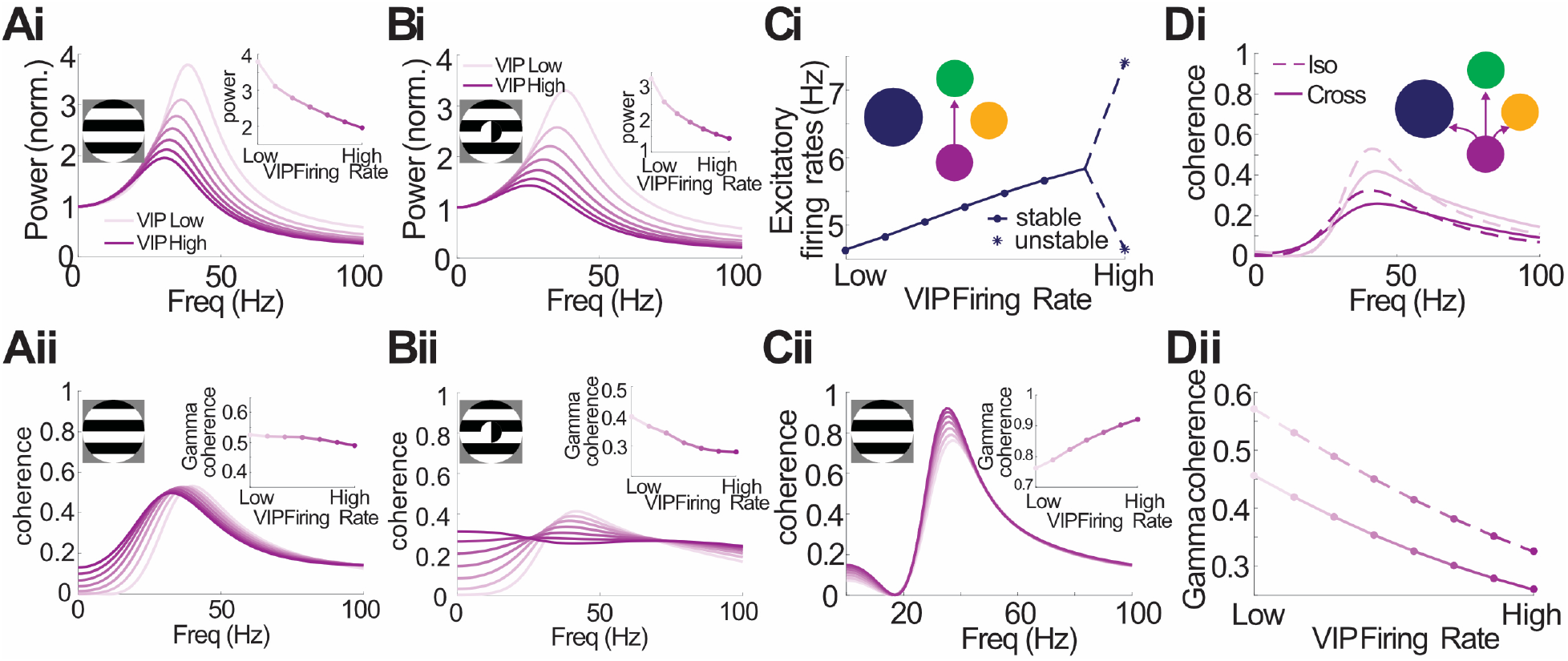
Connectivity of VIP neurons in the local circuit is crucial for controlling gamma coherence. **A:** Normalized gamma power (Ai) and coherence (Aii) for an iso-surround as a function of VIP firing rate. Despite gamma power decreasing as VIP firing rate increases (Ai, inset) the coherence at the gamma frequency remains relatively constant (Aii, inset). **B:** Same as A, except with a cross-surround. In this case, a decrease in gamma power coincides with a decrease in coherence at the gamma frequency. **C:** Simulations results for an iso-surround with a model where VIP inhibits PV, as opposed to SST (Ci, inset). Ci: Firing rate curve showing that stability is lost as VIP firing rates increase (dots are stable steady states, stars indicate the max and min of the oscillatory solution). Cii: Coherence curves, which, as opposed to the default model, shows an increase in gamma coherence as VIP firing rates increase (inset). **D:** Di: Coherence curves for an iso-(dashed) and cross-surround (solid) with high (dark lines) and low (light lines) VIP firing rates, where the model considers a VIP population that inhibits PC, PV, and SST populations with equal strength (inset). Dii: Gamma coherence decreases significantly for both the iso- and cross-surround conditions.

## Discussion

The data in this study establishes the disinhibitory VIP cell as a crucial regulator of gamma rhythms in the primary visual cortex. Importantly, optogenetically suppressing VIP neurons profoundly impacted the strength and spatial coherence of gamma rhythms, but did so in highly unexpected ways. Recent studies have highlighted the opposing responses of VIP and SST neurons to varied visual stimuli (Keller *et al.*, 2020; Millman *et al.*, 2020; Mossing *et al.*, 2021). While SST neurons are strongly driven by large high contrast, iso-oriented gratings, VIP neurons are suppressed by these stimuli and instead driven best by small, low contrast, or cross-oriented gratings. Two of these studies proposed or directly showed through optogenetic perturbations that VIP neurons tune the pyramidal network along these stimulus properties. All of this data supported a hypothesis wherein VIP neurons would likewise tune the stimulus-dependence of gamma oscillations. Strikingly and unexpectedly, our data refute this notion, as suppressing VIP neurons had a nearly exclusive impact on the gain of gamma rhythms, not their stimulus tuning. The most dramatic change in gamma power during VIP suppression was for the largest, and highest contrast iso-oriented gratings, while we observed fairly small effects for small, low contrast or cross-oriented gratings.

These data could have supported a relatively simple, albeit counterintuitive, model where the only role of VIP neurons in cortical gamma band rhythms was to control the gain of gamma band synchrony locally. However, our data with multi-site recording demonstrate that VIP neurons have a second, and arguably more important function in the global properties of gamma oscillations. VIP-mediated disinhibition, putatively through inhibition of SST neurons, preferentially suppressed inter-site coherence when the two sites were processing non-matched stimulus features, such as different orientations. Conversely, when these stimulus features matched, VIP neurons permit spatial coherence, even while they simultaneously scale down the total power of gamma divisively.

This raises a crucial next question: why do VIP neurons dampen gamma oscillations locally but permit coherence globally, depending on the visual context? One idea is that this local, divisive scaling of synchrony prevents the hyper-synchronization of cortical pyramidal neurons that might lead to aberrant propagation of activity to higher cortrical areas (Salinas & Sejnowski, 2001). In the same vein, VIP neuron activity might enhance visual perception by expanding the dynamic range of stimulus-dependent oscillatory dynamics. Importantly, even as they reduce local rhythmicity, VIP neurons allow distant network oscillators to couple when they are processing matched stimulus features. This might enhance the output of downstream neurons integrating across cortical space. VIP activity could thus act as a temopral filter in the gamma band for spatially continuous image features, such as contours and surfaces of objects.

The VIP-dependent decoupling between local and global neural synchronization argues that gamma power and coherence are not necessarily intrinsically linked. Superficially, this finding conflicts with a recent study which argues that gamma power and coherence should always be highly correlated (Schneider *et al.*, 2021). However, this study primarily considered sender and receiver populations that reside in different brain areas, rather that differentially tuned and spatially separate populations in the same cortical region, which we study here. Our theoretical work also differs from Schneider *et al.* (2021), in that we use standard techniques from non-equilibrium statistical mechanics to calculate the spike train power and cross-spectrums. As a result, a shift in the operating point (through changing contrast, VIP activation, cross vs. iso surround) changes the power and cross-spectrums in different ways. Since our formula for coherence depends on both of these quantities for a finite number of neurons, coherence will not be intrinsically linked to power without additional assumptions (see Supplemental Methods). In light of our experimental observations, such assumptions do not hold across spatial locations in V1.

A key outstanding question is what excitatory inputs drive VIP neurons, and in turn, how they mediate their divergent local and global effects. Although VIP neurons are known targets of corticocortical feedback axons from higher cortical areas, (Lee *et al.*, 2013; Zhang *et al.*, 2014) they are also local targets of V1 horizontal axons in layer 2/3 (Xu & Callaway, 2009; Karnani, Jackson, Ayzenshtat, Tucciarone, *et al.*, 2016). Our computational modeling implies that both the local and global action of VIP on gamma rhythms can be mediated entirely within V1. Analysis of the model revealed several key features that were important for the robustness of capturing the physiological results. First, and most intuitively, global coherence depended on specific like-to-like (i.e, iso-oriented) connectivity between center and surround circuits. Second, for capturing the selective effect of VIP on suppressing coherence to cross-oriented stimuli, it was important that VIP selectively targeted SST, as alternative models that generalized VIP neurons to target other cell types fell short in capturing our experimental results. Third, and perhaps least intuitively, in our model VIP neurons did not need tuned input from the local V1 network. Although VIP neurons do receive recurrent excitation from L2/3 PCs, our modeling surprisingly suggests that tuned excitatory input is not required for VIPs role in regulating the stimulus-dependence global coherence – rather they can enforce this effect through their powerful inhibition of SSTs which do get tuned recurrent input in the model.

Taken together, our data reveal a key new mechanism for the dynamic control of gamma-band neural synchronization in the primary visual cortex. As the same disinhibitory circuits exist in other sensory and higher cortical areas, the role of VIP neurons in controlling the gain and spatial coherence of gamma entrainment might be a general feature of cortical networks. Furthermore, our data suggest that VIP neurons might be potential therapeutic targets in neurological disorders that are associated with altered gamma rhythms and defects in inhibitory neural circuitry. Optogenetic or pharmacological tools aimed at re-balancing activity in VIP neurons, or perhaps more specific subsets of VIP neurons, should thus be useful in understanding the role of gamma rhythms in normal brain function and perhaps correcting it in disease.

---

**Figure S1.**
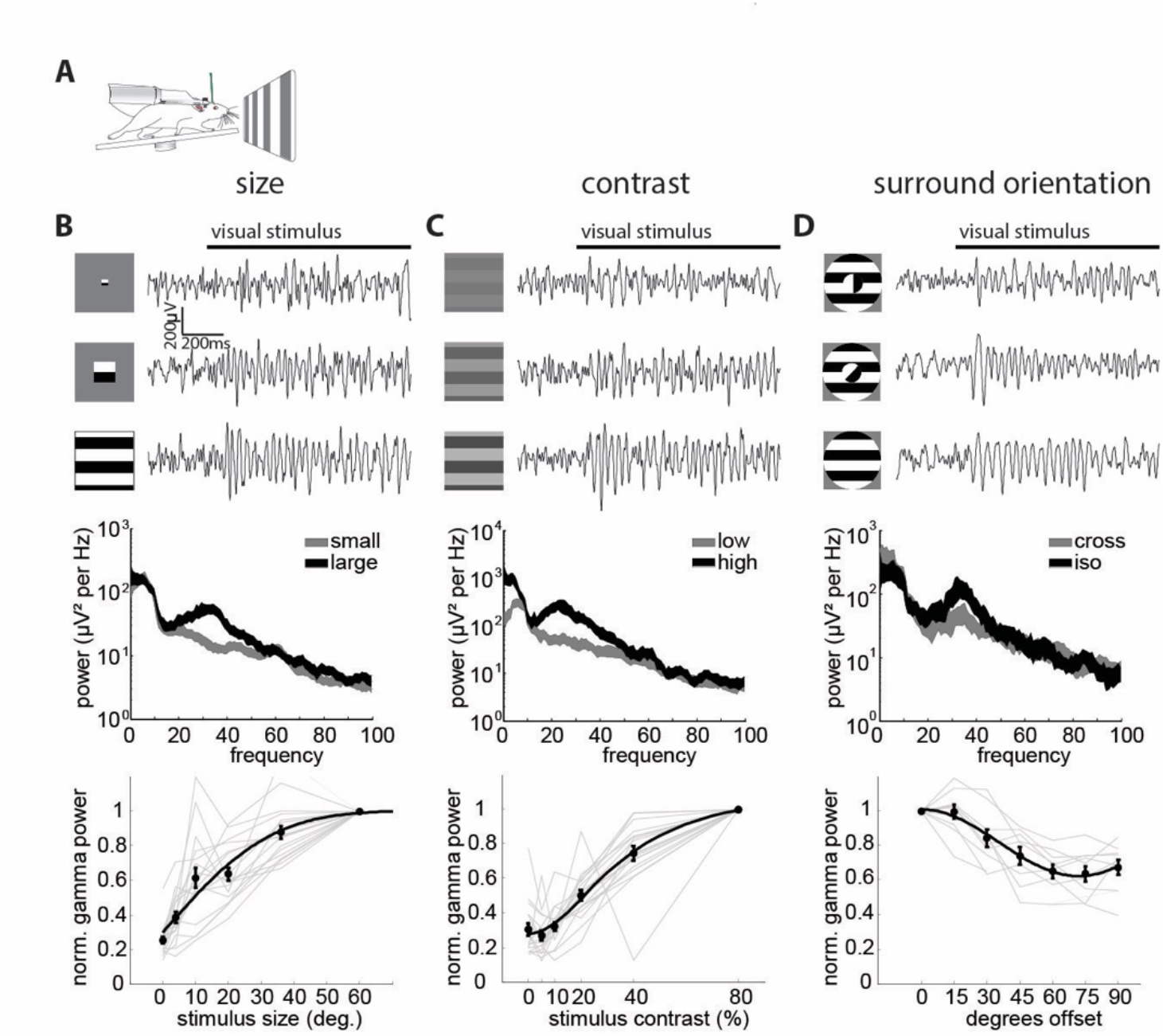
Stimulus dependence of the visually induced gamma rhythm in V1 of awake, running mice. **A:** Experimental schematic of a head-fixed mouse on a running wheel facing a screen for visual stimulation. **B:** Top: three example LFP traces filtered between 10 and 90 Hz in response to a small (4°, top), medium (20°, middle) and large (60°, bottom) drifting grating (temporal frequency, 2 Hz). Middle: example LFP power spectrum in response to a small (4**°**) and large (60**°**) drifting grating (thickness of line denotes mean ± standard error). Bottom: Plot of normalized peak gamma power (peak frequency: 28.8±0.5 Hz) versus stimulus size (Kruskal-Wallis ANOVA: p < 0.0001, n = 17) **C:** Top: three example filtered LFP traces in response to a low (5%, top), medium (20%, middle) and high (80%, bottom) contrast drifting grating. Middle: example LFP power spectrum in response to low (5%) and high (80%) contrast gratings (thickness of line denotes mean ± standard error). Bottom: Plot of normalized peak gamma power (peak frequency: 27.3±0.6 Hz) versus stimulus contrast (Kruskal-Wallis ANOVA: p<0.0001, n = 18). **D:** Top: three example filtered LFP traces in response to a grating with cross-(90° offset, top), intermediate-(45° offset, middle) and iso-oriented (0° offset, bottom) surround relative to the center. Middle: Example LFP power spectrum in response to iso and cross oriented surround grating (thickness of line denotes mean ± standard error). Bottom: Plot of normalized peak gamma power (peak frequency: 32.8±0.4 Hz) versus stimulus contrast (Kruskal-Wallis ANOVA: p<0.0001, n = 10). Error bars in all plots represent s.e.m.

**Figure S2:**
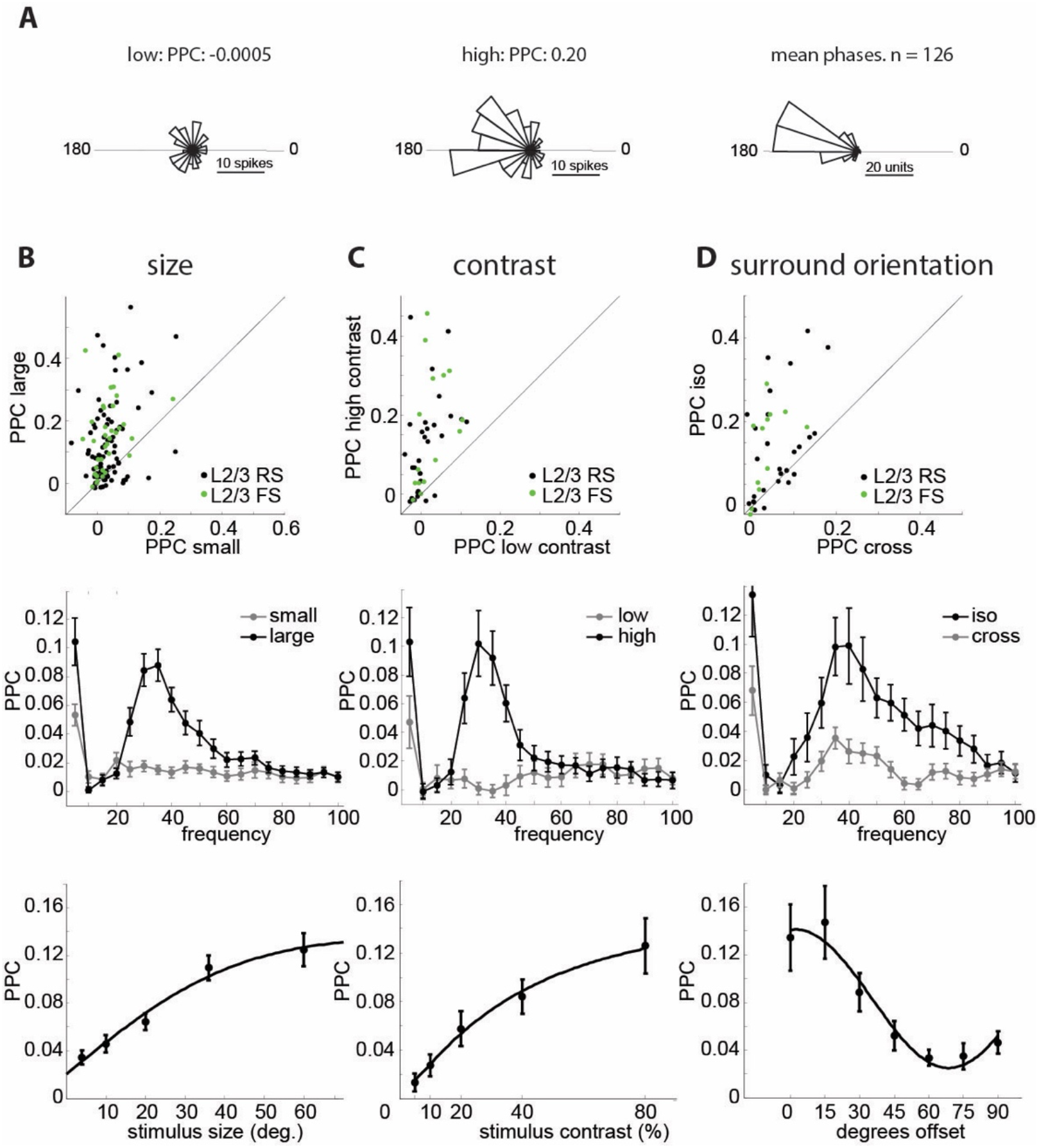
Single units lock to visually induced gamma oscillations in a stimulus-dependent manner. **A:** Left: phase histogram of the spikes of an example L2/3 RS unit relative to the gamma oscillation in response to a low contrast (5%) stimulus. Middle: similar histogram from the spikes of the same neuron evoked by a high contrast (80%) stimulus. Right: histogram of the average spike phases of all 126 included L2/3 RS cells included for a large high contrast grating. Cells tend to fire shortly before the trough of the oscillation (180°). **B:** Top: scatter plot of PPC values for single RS (black, n = 78, p<0.0001, Wilcoxon signed rank test) and FS (green, n = 32, p<0.0001, Wilcoxon signed rank test) units in response to small (4°) and large (60°) full contrast stimuli. Middle: average PPC spectra for L2/3 RS cells (n = 78) for small (gray, 4°) and large (black, 60°) full contrast stimuli. Bottom: Plot of average PPC at individual gamma center frequency versus stimulus size for L2/3 RS units (n = 87, p < 0.0001, Kruskal-Wallis ANOVA). **C:** Top: scatter plot of PPC values for single RS (black, n = 29, p<0.0001, Wilcoxon signed rank test) and FS (green, n = 15, p<0.0001, Wilcoxon signed rank test) units in response to large low (5%) and high (80%) contrast stimuli. Middle: average PPC spectra for L2/3 RS cells (n = 29) for large low (gray, 5%) and high (black, 80%) contrast stimuli. Bottom: plot of average PPC at individual gamma center frequency versus stimulus contrast for L2/3 RS units (n = 29, p < 0.0001, Kruskal-Wallis ANOVA). **D:** Top: scatter plot of PPC values for single RS (black, n = 27, p = 0.0008, Wilcoxon signed rank test) and FS (green, n = 13, p = 0.003, Wilcoxon signed rank test) cells in response to full contrast cross (90° offset) and iso (0° offset) surround stimuli. Middle: average PPC spectra for L2/3 RS cells (n = 27) for full contrast cross (gray, 90° offset) and iso (black, 0° offset) surround stimuli. Bottom: plot of average PPC at individual gamma center frequency versus relative surround orientation for L2/3 RS units (n = 28, p = 0.004, Kruskal-Wallis ANOVA). Error bars in all plots represent s.e.m.; see Supp Fig 1 for FS data.

**Figure S3:**
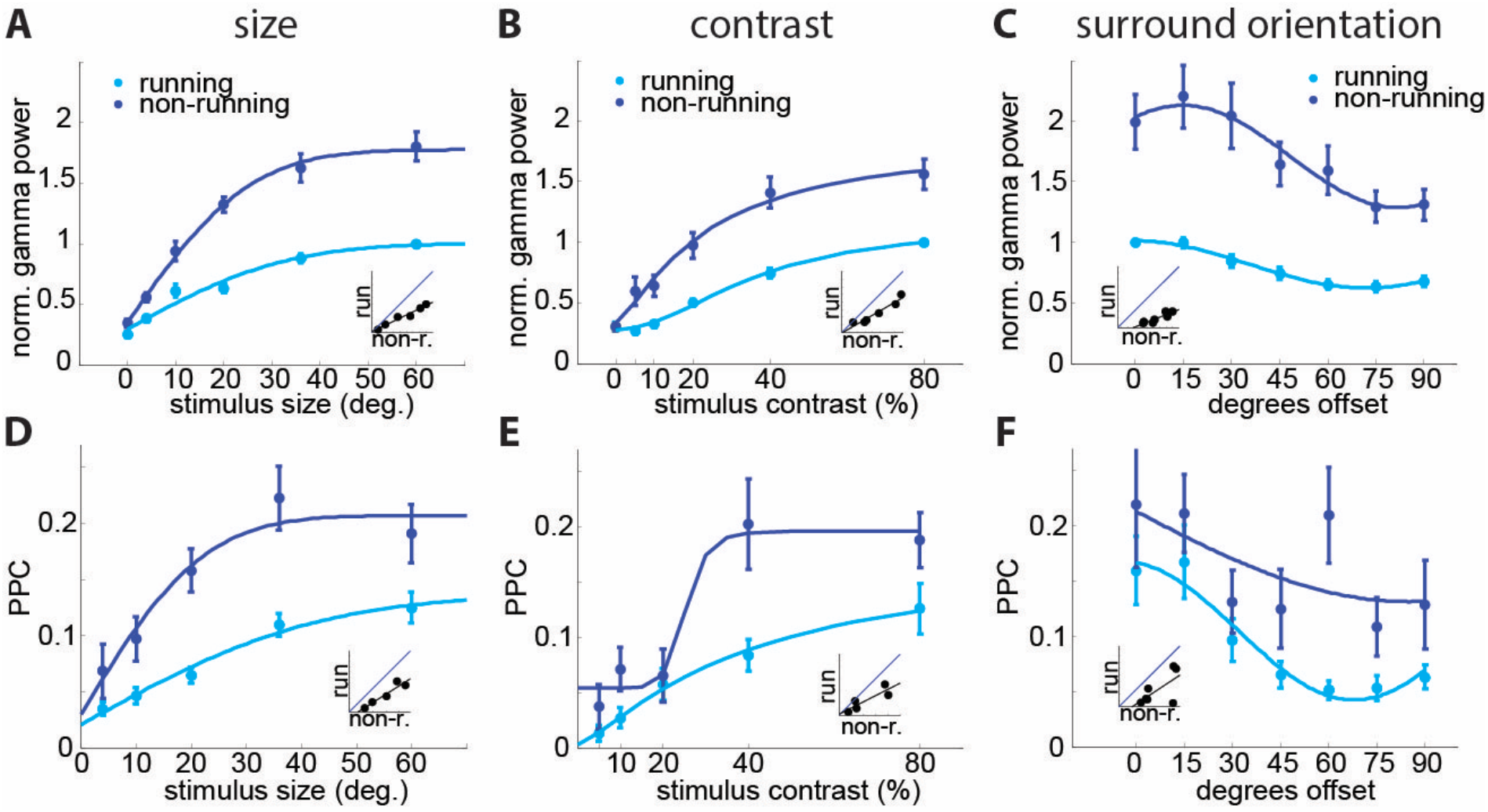
Gamma power and phase locking depend on behavioral state. **A:** Average normalized gamma power during running (light blue) and non-running (dark blue) versus stimulus size (n = 17, 2-way-ANOVA: main effect of size: F(4,146) = 45.34, p<0.001; main effect of running F(1,146) = 122.49, p<0.001; interaction: F(4,146) = 6.40, p<0.001). **B:** Average normalized gamma power during running and non-running versus stimulus contrast (n = 18, 2-way ANOVA: main effect of contrast: F(4,149) = 33.68, p<0.001; main effect of running: F(1,149) = 67.82, p<0.001; interaction: F(4,149) = 1.39, p = 0.24). **C:** Average normalized gamma power during running and non-running versus relative surround orientation (n = 10, 2-way ANOVA: main effect of orientation: F(6,108) = 6.38, p<0.001; main effect of running: F(1,108) = 156.02, p<0.001; interaction: F(6,108) = 1.34, p = 0.24) **D:** Average PPC during running (light blue) and non-running (dark blue) versus stimulus size (n = 87, 2-way-ANOVA: main effect of size: F(4,835) = 16.3, p<0.001; main effect of running: F(1,835) = 37.74, p<0.001; interaction F(4,835): 1.1, p = 0.36). **E:** Average PPC during running and non-running versus stimulus contrast (n = 29, 2-way-ANOVA: main effect of contrast: F(4,256) = 14.02, p<0.001; main effect of running: F(1,256) = 13.49, p<0.001; interaction: F(4,256) = 1.9, p = 0.11). **F:** Average PPC during running and non-running versus relative surround orientation (n = 28, 2-way-ANOVA: main effect of orientation: F(6,328) = 3.75, p= 0.001; main effect of running: F(1,328) = 15.46, p<0.001; interaction: F(6,328) = 0.84, p = 0.54). Error bars in all plots represent s.e.m.

**Figure S4:**
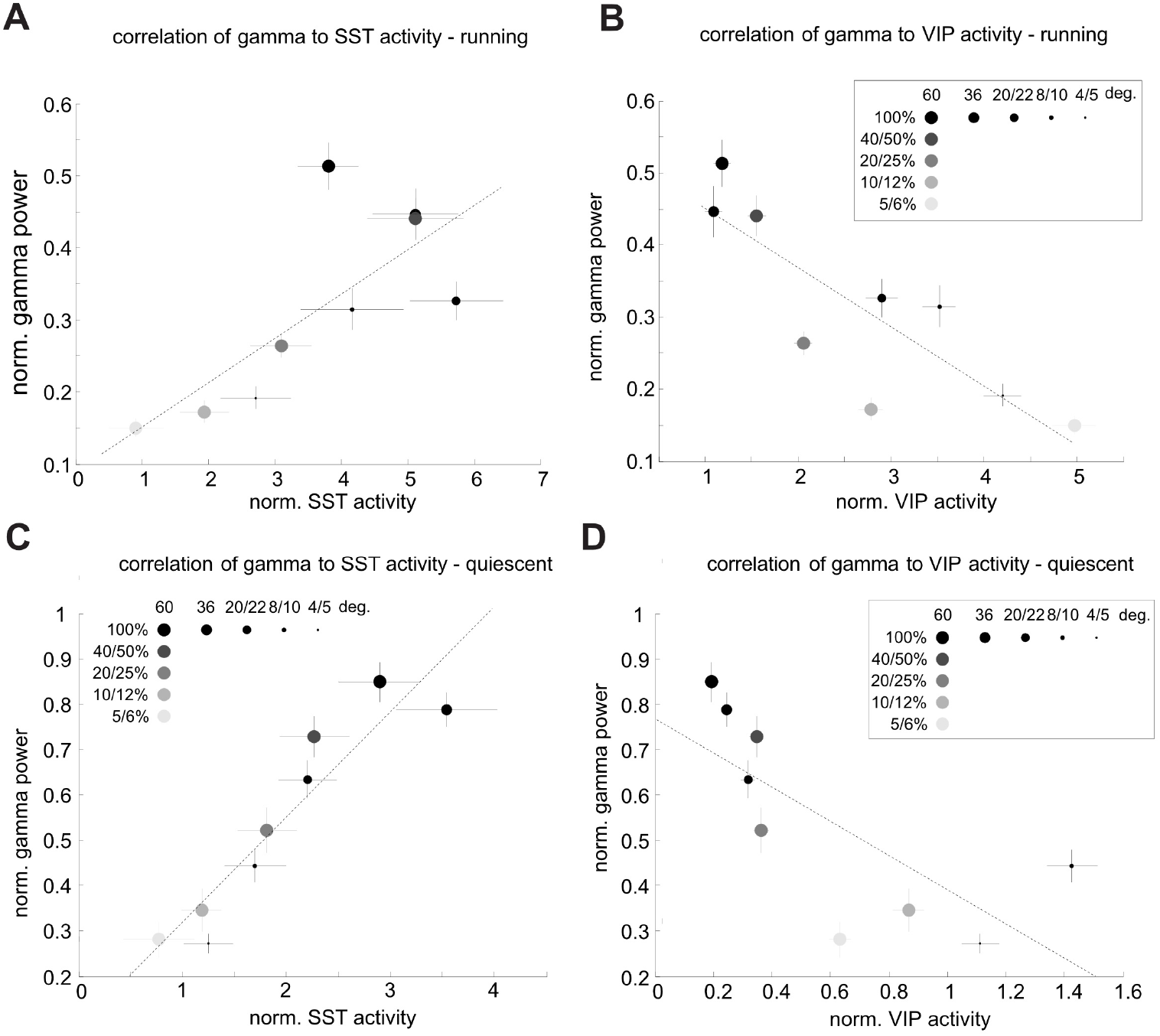
Opposing correlation of SST- and VIP-neuron activity with gamma power. **A:** Plot of averaged normalized gamma power in the running condition vs. averaged normalized SST-cell activity (deconvolved event-rate/mean), recorded via 2-photon imaging in a different set of animals across similar conditions. Different shades of gray represent different contrast levels and different symbol sizes represent different stimulus sizes. Dashed line is a linear fit of the data. SST-cell activity strongly correlates with gamma power (r(7) = 0.76, p = 0.019). **B:** Same as A, except for normalized VIP cell activity. VIP activity is strongly anti-correlated with gamma power (r(7) = −0.84, p = 0.005). **C:** Same as A, except in the quiescent state. SST-cell activity is strongly correlated to gamma power (r(7) = 0.93, p<0.001). **B:** Same as B, except in the quiescent state. VIP activity is strongly anti-correlated to gamma power (r(7) = −0.73, p = 0.024).

**Figure S5:**
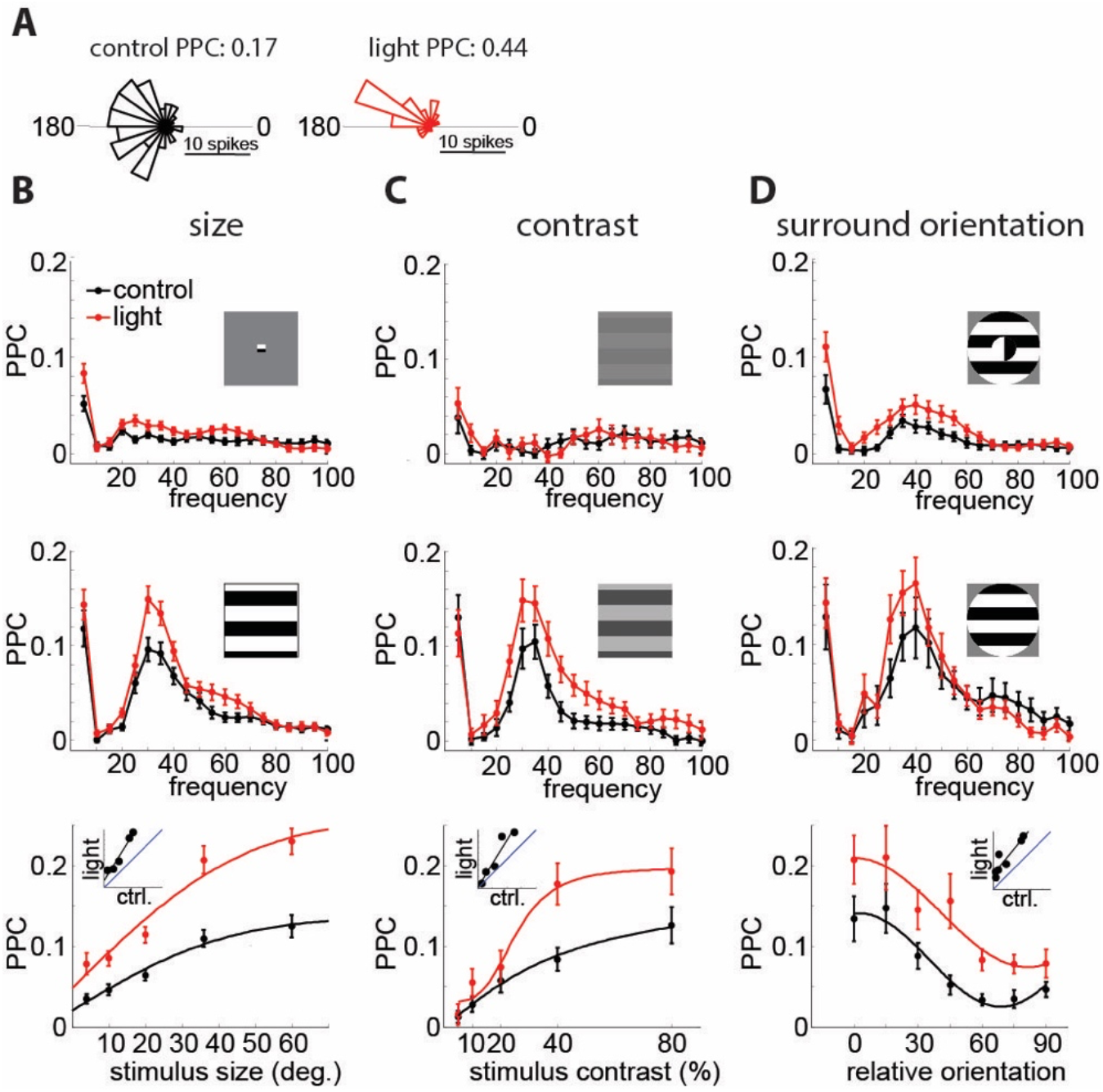
VIP neurons linearly control the strength of locking of single neurons to the visually induced gamma rhythm. **A:** Left: phase histogram of the spikes of an example L2/3 RS neuron relative to the gamma oscillation in the control condition (60° grating, same unit as in Fig. 2). Right: phase histogram of the spikes of the same neuron during inactivation of VIP neurons. **B:** Top: average PPC spectra for L2/3 RS cells with (red) and without (black) suppression of VIP neurons (n = 78) for small (4°) stimuli. Middle: average PPC spectra for L2/3 RS cells with (red) and without (black) suppression of VIP neurons (n = 68) for large (60°) stimuli. Bottom: Plot of average PPC versus stimulus size with (red) and without (black) light-mediated inactivation of VIP neurons (n = 87, 2-way ANOVA: main effect of light: F(1,857) = 78.42, p<0.001; main effect of size: F(4,857) = 42.83, p<0.001; interaction: F(4,857) = 3.14, p = 0.014). **C:** Top: average PPC spectra for L2/3 RS cells with (red) and without (black) suppression of VIP neurons (n = 27) for low contrast (5%) stimuli. Middle: average PPC spectra for L2/3 RS cells with (red) and without (black) suppression of VIP neurons (n = 30) for high contrast (80%) stimuli. Bottom: Plot of average PPC versus stimulus contrast with (red) and without (black) inactivation of PV neurons (n = 29, 2-way ANOVA: main effect of light: F(1,280) = 13.01, p<0.001; main effect of contrast: F(4,280) = 21.8, p<0.001; interaction: F(4,280) = 2.17, p = 0.072). **D:** Top: average PPC spectra for L2/3 RS cells with (red) and without (black) suppression of VIP neurons (n = 46) for cross surround (90° offset) stimuli. Middle: average PPC spectra for L2/3 RS cells with (red) and without (black) suppression of VIP neurons (n = 21) for iso surround (0° offset) stimuli. Bottom: Plot of average PPC versus relative surround orientation with (red) and without (black) inactivation of VIP neurons (n = 28, 2-way ANOVA: main effect of light: F(1,378) = 28.66, p<0.001; main effect of orientation: F(6,378) = 10.31, p<0.001; interaction: F(6,378) = 0.69, p = 0.66). Error bars in all plots represent s.e.m.

**Fig. S6.**
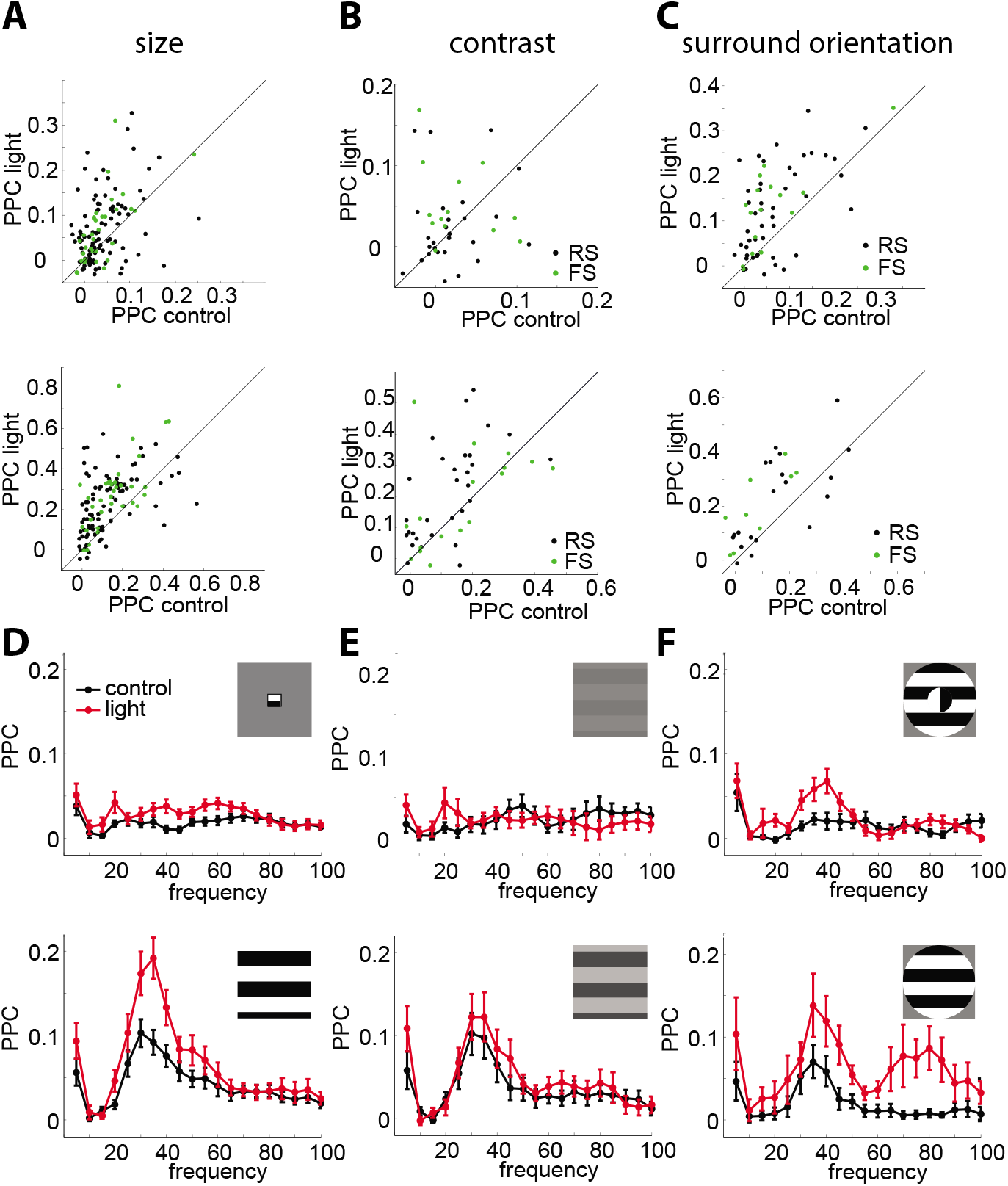
Effects of VIP inactivation on locking of single RS and FS cells. **A:** Top: scatter plot of PPC values for single RS (black, n = 90, p = 0.0001, Wilcoxon signed rank test) and FS (green, n = 33, p = 0.002, Wilcoxon signed rank test) cells in response to small (4°) stimuli in control condition versus VIP suppression. Bottom: scatter plot of PPC values for single RS (black, n = 87, p = 0.0004, Wilcoxon signed rank test) and FS (green, n = 35, p = 0.002, Wilcoxon signed rank test) cells in response to large (60°) stimuli in control condition versus VIP suppression. **B:** Top: scatter plot of PPC values for single RS (black, n = 27, p = 0.61, Wilcoxon signed rank test) and FS (green, n = 13, p = 0.31, Wilcoxon signed rank test) cells in response to low contrast (5%) stimuli in control condition versus VIP suppression. Bottom: scatter plot of PPC values for single RS (black, n = 30, p = 0.001, Wilcoxon signed rank test) and FS (green, n = 17, p = 0.98, Wilcoxon signed rank test) cells in response to large (60°) stimuli in control condition versus VIP suppression. **C:** Top: scatter plot of PPC values for single RS (black, n = 46, p < 0.0001, Wilcoxon signed rank test) and FS (green, n = 15, p = 0.0006, Wilcoxon signed rank test) cells in response to cross surround stimuli in control condition versus VIP suppression.Bottom: scatter plot of PPC values for single RS (black, n = 21, p = 0.04, Wilcoxon signed rank test) and FS (green, n = 9, p = 0.004, Wilcoxon signed rank test) cells in response to iso surround (0° offset) stimuli in control condition versus VIP suppression. **D:** Top: average PPC spectra for L2/3 FS cells with (red) and without (black) suppression of VIP neurons (n = 30) for small (4°) stimuli. Bottom: average PPC spectra for L2/3 FS cells with (red) and without (black) suppression of VIP neurons (n = 32) for large (60°) stimuli. **E:** Top: average PPC spectra for L2/3 FS cells with (red) and without (black) suppression of VIP neurons (n = 13) for low contrast (5%) stimuli. Bottom: average PPC spectra for L2/3 FS cells with (red) and without (black) suppression of VIP neurons (n = 17) for high contrast (80%) stimuli. **F:** top: average PPC spectra for L2/3 FS cells with (red) and without (black) suppression of VIP neurons (n = 15) for cross surround (90° offset) stimuli. Bottom: average PPC spectra for L2/3 FS cells with (red) and without (black) suppression of VIP neurons (n = 9) for iso surround (0° offset) stimuli.

**Figure S7.**
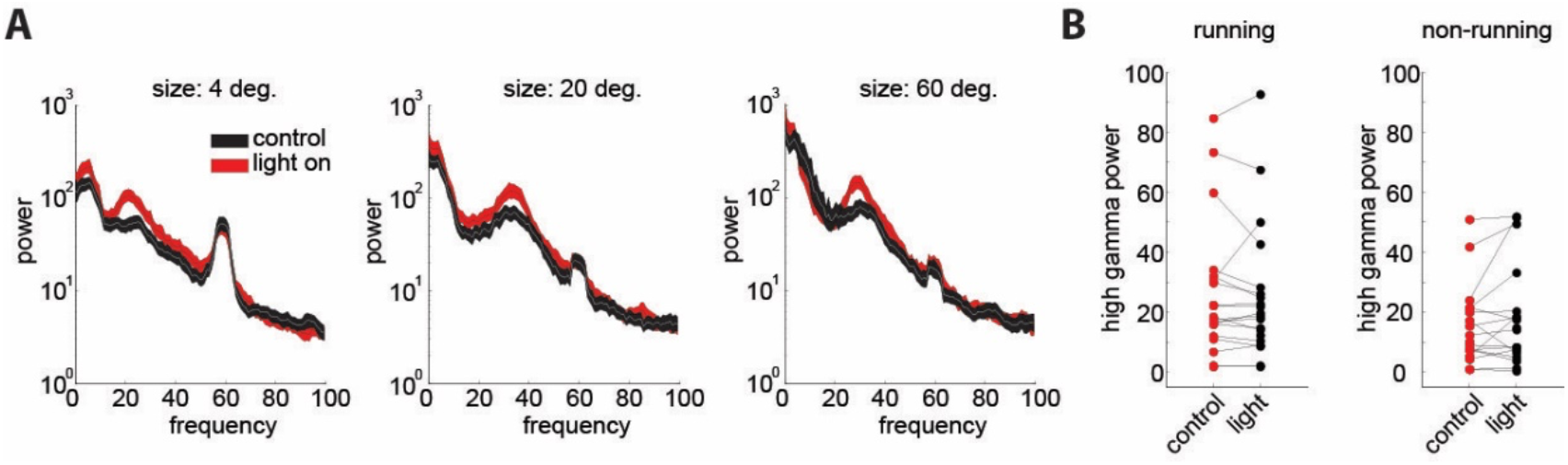
Effects of VIP inactivation on higher-frequency, narrowband, thalamic gamma (60Hz). **A:** Spectra for different size grating stimuli with (red) and without (black) inactivation of VIP neurons. VIP affects the visually induced 30Hz gamma band, but not the thalamically relayed 60Hz gamma band that is supressed by large/high contrast stimuli. **B:** Plot comparing the LFP high gamma band power for blank stimuli in the running condition for control (black) and light (red) trials (n = 19, p = 0.33, Wilcoxon signed rank test) Right: Plot comparing the LFP high gamma band power for blank stimuli in the non-running condition for control (black) and light (red) trials (n = 18, p = 0.25, Wilcoxon signed rank test).

**Figure S8:**
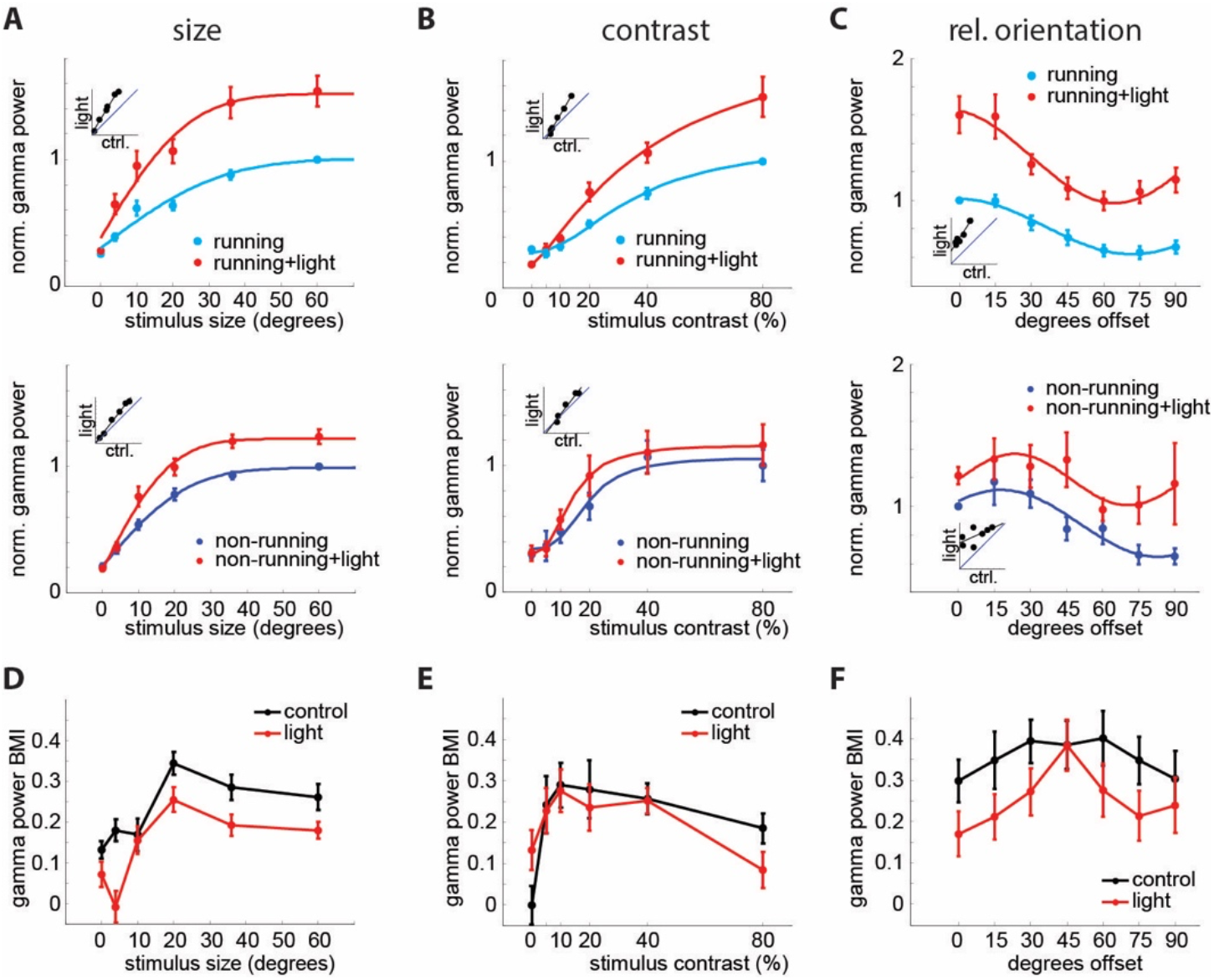
VIP neurons contribute to the behavioral dependence of gamma band synchronization. **A:** Top: Average normalized gamma power as a function of stimulus size with (red) and without optogenetic suppression of VIP neurons during running (light blue, n = 21, 2-way ANOVA: mian effect of light: F(1,160) = 54.18, p<0.001; main effect of size: F(4,160) = 22.18, p<0.001; interaction: F(4,160) = 1.03, p = 0.39) Bottom: Same for non-running (dark blue, n = 21, 2-way-ANOVA: main effect of light: F(1,130) = 26.49, p<0.001; main effect of size: F(4,130) = 55.9, p<0.001; interaction: F(4,130) = 1.41, p = 0.23). **B:** Top: Average normalized gamma power as a function of stimulus contrast with (red) and without optogenetic suppression of VIP neurons during running (light blue, n = 21, 2-way ANOVA: main effect of light: F(1,170) = 27.81, p<0.001; main effect of contrast: F(4,170) = 65.08, p<0.001; interaction: F(4,170) = 3.85, p = 0.005) Bottom: Same for non-running (dark blue, n = 21, 2-way-ANOVA: main effect of light: F(1,125) = 1.48, p = 0.23; main effect of contrast: F(4,125) = 12.36, p<0.001; interaction: F(4,125) = 0.31, p = 0.87). **C:** Top: Average normalized gamma power as a function of relative surround angle with (red) and without optogenetic suppression of VIP neurons during running (light blue, n = 10, 2-way ANOVA: main effect of light: F(1,125) = 119.37, p<0.001; main effect of orientation: F(6,125) = 13.14, p<0.001; interaction: F(6,125) = 0.88, p = 0.51) Bottom: Same for non-running (dark blue, n = 7, 2-way-ANOVA: main effect of light: F(1,83) = 11.27, p= 0.001; main effect of orientation: F(6,83) = 1.9, p = 0.09; interaction: F(6,83) = 0.47, p = 0.83). **D:** Average gamma power BMI (behavioral modulation index) as a function of stimulus size with (red) and without (black) optogenetic suppression of VIP neurons (n = 21, 2-way-ANOVA: main effect of light: F(1,134) = 14.34, p<0.001; main effect of size: F(4,134) = 8.84, p<0.001; interaction: F(4,134) = 1.29, p = 0.28). **E:** Average gamma power BMI as a function of stimulus contrast with (red) and without (black) optogenetic suppression of VIP neurons (n = 21, 2-way-ANOVA: main effect of light: F(1,130) = 0.66, p = 0.42; main effect of contrast: F(4,130) = 1.79, p = 0.14; interaction: F(4,130) = 0.22, p = 0.93). **F:** Average gamma power BMI as a function of relative surround angle with (red) and without (black) optogenetic suppression of VIP neurons (n = 10, 2-way-ANOVA: main effect of light: F(1,92) = 7.44, p = 0.008; main effect of orientation: F(6,92) = 1.03, p = 0.41; interaction: F(6,92), p = 0.96).

**Figure S9.**
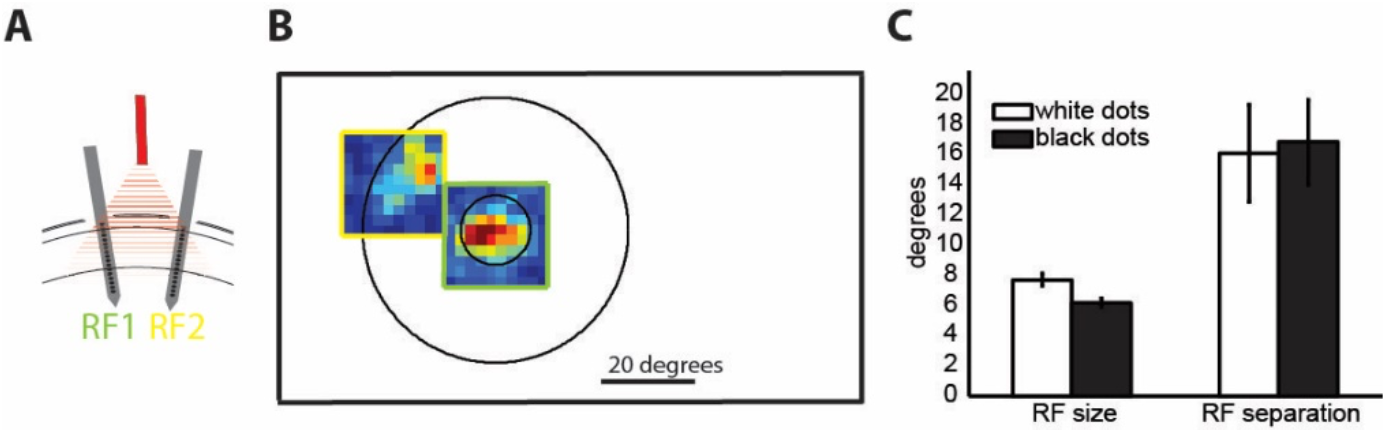
Receptive field mapping procedure for coherence measurement. **A:** Schematic of the multielectrode array recording configuration with two laminar arrays in distant sites (530+-90 μm apart, histology from n = 7 mice) corresponding to two separate retinotopic locations (RF1 (green) and RF2 (yellow), 15° ± 3° of visual angle separation, n = 11 mice). Red triangle denotes wide illumination with optogenetic light delivered from a fiber located above the two recording sites. **B:** Two sparse noise mapped RFs (redder colors denote higher firing rates), one from electrode 1 (green frame), one from electrode 2 (yellow frame) superimposed on the outline of the center and surround of the visual stimulus used for the coherence analysis (Figure 7). Large outer frame is approximately the size of the stimulation monitor. **C:** average RF size (2 standard deviations of Gaussian fit to RF) and average separation of center and surround fields, separately for fields mapped with white and black sparse noise, n = 8.

## Materials and Methods

### Transgenic mice

All experiments were performed in accordance with the guidelines and regulations of the ACUC of the University of California, Berkeley. Mice for the *in vivo* experiments were housed in groups of five or less with a 12:12h light:dark cycle. Both female and male mice were used. Experiments *in vivo* were performed on animals aged between 8–27 weeks during their subjective night. We used VIP-IRES-Cre (JAX stock 010908) mice. Mice were out-crossed for one generation to the ICR white strain (Charles River).

### Viral infection

Neonatal VIP-Cre mice (P3–6) were briefly cryo-anesthetized and placed in a head mold. Transcranial injection of ~45nl of undiluted AAV9-EF1a-DIO-eNpHR3.0-YFP (22 animals) was performed using a Drummond Nanoject injector at three locations in V1 using a glass pipette beveled to fine tip (~30-60μm). With respect to the lambda suture coordinates for V1 were 0.0 mm AP, 2.2 mm L and injection was as superficial as possible under the skull.

### *Preparation for* in vivo *recording*

Mice were anesthetized with isoflurane (2.5% vapor concentration). The scalp was removed, the fascia retracted, and the skull lightly etched with a 27 gauge needle. Following application of Vetbond to the skull surface, a custom stainless steel headplate was fixed to the skull with dental cement (Metabond). Mice were allowed to recover from surgery for at least 2 days. Then mice were habituated for 2–10 days to head-fixation on a free-spinning circular treadmill. On the day of recording mice were briefly anesthetized with isoflurane (2%), the skull over V1 was thinned, and one or two (spacing 400-1000μm) small (<250 μm) craniotomies were opened over V1 with a fine needle

### Visual stimulation

Visual stimuli were generated with Psychophysics Toolbox (Brainard, 1997) running on an Apple Mac Mini and were presented on a gamma corrected 23-inch Eizo FORIS FS2333 LCD display with a 60-Hz refresh rate. At the beginning of each recording session the receptive fields of MUA recorded at each cortical location was mapped with sparse noise to be able to precisely position the grating stimuli. The stimulus was centered on a location where a small grating, movable by hand, elicited a clear response. Sparse noise consisted of black and white squares (2 visual degrees, 80 ms) on a 20×20 visual degree grid flashed onto a gray background of intermediate luminance. To improve receptive field estimation the same stimulus grid was offset by 1 degree and the resulting maps were averaged. MUA average receptive fields were calculated by reverse correlation.

Visual stimuli consisted of drifting square-wave gratings at 0.04 cycles per degree and 2 cycles per second centered on the average MUA receptive field presented for 2s with at least 1s inter stimulus interval. Gratings were presented in three different configurations: 1) full contrast gratings of eight different directions (0–315° in steps of 45°) and five different sizes (4 10, 20, 36, and, if possible, 60 visual degrees – if the RF was not perfectly centered on the monitor, the effective largest size was slightly smaller); 2) gratings of four different directions (0-270° in steps of 90°), three different sizes (8,20 and 60°) and 5 different contrast levels (0.05, 0.1, 0.2, 0.4, 0.8) Michelson contrast and 3) full contrast square-wave gratings with a circular aperture of 8-15° visual degrees diameter (depending on the separation of the two RFs), centered on the MUA receptive field of one of the two simultaneously recorded cortical locations, that was surrounded by a 60 degree grating with one of seven different relative orientations (0-180° in steps of 30°). For the coherence analysis we only analyzed cases in which the second receptive field was covered entirely and exclusively by the surround-stimulus (see Fig. 7A and sup. Fig. 2).

### Optogenetic stimulation in vivo

For optogenetic stimulation of eNpHR3.0 in vivo we used red (center wavelength: 625 nm) from the end of a 1-mm diameter multimode optical fiber coupled to a fiber coupled LED (Thorlabs) controlled by digital outputs (NI PCIe-6353). The fiber was placed as close to the craniotomy as possible (<3 mm). The illumination area was set to illuminate a wide area including all of V1. Light levels were tested in increasing intensities at the beginning of the experiment and were kept at the lowest possible level that still evoked observable change in ongoing activity for the remainder of the recording. We only used viral injections into V1, and did not attempt to use an eNphR transgenic reporter line to avoid off-target expression of the opsin and non-specific optogenetic suppression of subcortical nuclei (such as the thalamic reticular nucleus).

Gratings drifted for 2s with at least 1s inter-trial intervals with the red LED switched on for 1 s starting 0.5 s after start of the visual stimulus in 50% of the trials. The period of light was chosen to influence the stable steady-state of the response to the grating and all analysis was performed during this time window.

### In vivo extracellular multi-electrode electrophysiology

One or two 16-channel linear electrodes with 25 micron spacing (NeuroNexus, A1×16-5mm-25-177-A16) were guided into the brain using micromanipulators (Sensapex) and a stereomicroscope (Leica). Electrical activity was amplified and digitized at 30 kHz (Spike Gadgets), and stored on a computer hard drive. The cortical depth of each electrical contact was determined by zeroing the bottom contact to the surface of the brain. Electrodes were inserted close to perpendicular to the brain’s surface for single electrode recordings and ~25 degrees from vertical for the two electrode experiments. After each recording a laminar probe coated with the lipophilic dye DiI was used to mark each electrode track to quantitatively assess insertion angle and depth with post-hoc histologic reconstructions. The laminar depth of recorded units was corrected for the insertion angle and the local curvature of the neocortex.

### Analysis of local field potential data

All analysis was performed using custom written code or openly available packages in Matlab (Mathworks). Local field potentials were extracted by low pass filtering the raw signal, sampled at 30 kHz, below 200 Hz and subsequent down-sampling to 1 kHz. For LFP-only analysis we always analyzed the LFP from the electrode contact closest to a cortical depth of ~350 μm (in cortical layer 3). For spike locking to the LFP we used the LFP from an electrode contact 50 μm away from the contact with the largest spike-waveform amplitude to reduce contamination of the LFP.

The power spectrum was computed in a 800 ms analysis window starting 200 ms after light onset (to exclude any photo-electric artifacts sometimes present in the first ~150 ms after light onset) using multi-taper estimation in Matlab with the Chronux package (http://chronux.org/, Mitra & Bokil, 2007) using 3 tapers. All power analysis was performed on the power at the peak of each animal’s specific gamma oscillation in the specific visual stimulation condition. Peaks were identified as local maxima on the smoothed spectrum between 20 and 40Hz that were preceded by local minima in the 15Hz preceding the peak. If no true peak could be found (as was often the case for very small or low contrast conditions), we took the power at the frequency of the peak for the highest contrast/largest stimulus of that animal.

For calculation of coherence, bipolar derivatives of the LFP were calculated by subtracting the electrode channel two contacts above the channel of interest (50μm distance), to remove the common recording reference and to enhance spatial specificity of the signal. Coherence between the two recording sites was determined using the chronux package with the same number of tapers as the power analysis. All spectral plots show mean±s.e.m, the coherence spectra show jack-knifed 95% confidence intervals. Coherence values for the analysis were taken of the peak of each animals’ individual coherence spectrum as for the power above.

### Analysis of spiking data

Spiking activity was extracted by filtering the raw signal between 800 and 7000 Hz. Spike detection was performed using the UltraMega Sort package (Hill et al., 2011). Detected spike waveforms were sorted using the MClust package (http://redishlab.neuroscience.umn.edu/MClust/MClust.html). Waveforms were first clustered automatically using KlustaKwik and then manually corrected to meet criteria for further analysis. With the exception of <25 burst firing units, included units had no more than 1.5% of their individual waveforms violating a refractory period of 2 ms. Individual units were classified as either fast-spiking or regular spiking using a k-means cluster analysis of spike waveform components. Since the best separation criterion was the trough-to-peak latency of the large negative going deflection and clustering is non-deterministic, we defined all units with latencies shorter than 0.36 ms as fast spiking and all units with latencies larger than 0.38ms as regular spiking. Cells with intermediate latencies were excluded from further analysis.

The depth of each unit was assigned based on the calculated depth of the electrode on the array that exhibited its largest amplitude sorted waveform. Layer boundaries were determined following a previously established approach (Pluta *et al.*, 2015). Firing rates were computed from counting spikes in a 1 second window starting 500 ms after the onset of the visual stimulus, which coincided with the onset of the LED during optogenetic suppression trials. Unless otherwise stated, we only analyzed trials when the animal was moving (at least 1cm/s) and not accelerating or decelerating abruptly (not more than 1.5 s.d. deviation from the animal’s mean running speed).

To quantify locking of spiking activity to the gamma band we bandpass filtered the LFP in a 20 Hz band around the individual gamma band peak (between 20 and 45 Hz) and extracted the oscillation’s instantaneous phase by using the imaginary part of the analytical signal using the Hilbert transform. Each spike is thus assigned an exact phase in the gamma oscillation. Phase locking magnitude is determined for each unit by the pairwise phase consistency (PPC), a measure of synchrony that is not biased by the number of spikes (Vinck *et al.*, 2010). We only included units that fired more than 20 spikes total in response to the largest grating size in the control condition and whose average visual response rate was >1Hz. PPC-spectra were calculated as above but for LFP filtered into 20 non-overlapping 5Hz wide frequency bands.

Behavioral modulation index (BMI) was calculated as 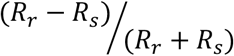 where *R*_*r*_ is the average response during running and *R*_*s*_ is the average response in non-running (still) trials.

For illustrative purposes the average functions for gamma power and PPC were fit with functions. For size tuning curves an integral of Gaussian, for contrast tuning a Naka-Rushton function and for center-surround angle a sinusoid was fit with Matlab curve fitting toolbox.

### Imaging data

Imaging data was performed as described in (Mossing *et al.*, 2021). Briefly, Sst-IRES-Cre and Vip-IRES-Cre mice wer crossed to Ai162(TIT2L-GC6s-ICL-tTA2)-D mice (RRID:IMSR_JAX:031562) and an imaging window implanted. The visual stimulus consisted of square wave drifting gratings, with directions tiling 0-360 degrees at 45° intervals, with a spatial frequency of 0.08 cycles per degree, and a temporal frequency of 1 Hz. Visual stimulus presentation lasted one second,followed by a one second inter-stimulus interval. Mice were head-fixed on a freely spinning running wheel under a Nixon 16x-magnification water immersion objective and imaged with a two-photon resonant scanning microscope (Neurolabware) within a light tight box. The imaging FOV was 430 by 670 um, with four planes spaced 37.5 μm apart imaged sequentiallyusing an electrotunable lens (Optotune), sampling each plane at an effective frame rate of 7.72 Hz. Motion correction and ROI segmentation was performed using Suite2p (Pachitariu *et al.*, 2017). Neuropil subtraction was applied as described in (Pluta *et al.*, 2017). ΔF/F traces were calculated as 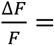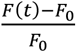 with baseline *F*_0_ computed over a sliding 20^th^ percentile filter of width 3000 frames. Because the inter-stimulus interval was short to permit more stimuli to be displayed, calcium transients overlapped between successive trials. Therefore, we deconvolved calcium traces for this data using OASIS with L1 sparsity penalty (Friedrich *et al.*, 2017) using ΔF/F traces as input. We report this deconvolved event rate normalized by the mean.

### Mathematical methods

We modeled a network of exponential integrate-and-fire neurons. Simulations were completed using Euler’s method using a timestep of 0.025 msec for a total of 1e6 msec of simulation time. The auto- and cross-correlation functions were then estimated by binning the spike times over 1 msec time windows, summing this count across excitatory neurons in a retinotopic location, and then using MATLAB’s built-in xcorr() function for a max window length of 250 msec. The power spectrum and cross-spectrum were then computed by taking the Fourier transform. Additional descriptions of the spiking model, linear response theory and mean-field model can be found in the Supplemental Methods. The code to reproduce the key figures from the computational model can be found on GitHub (https://github.com/gregoryhandy).

## Acknowledgements

This work was funded by the New York Stem Cell Foundation. H.A. is a New York Stem Cell Foundation Robertson Investigator. This work was supported by NEI grant R01EY023756 and NINDS grant U19NS107613. B.D. was supported by NIH grants 1U19NS107613-01 and R01 EB026953, the Vannevar Bush Faculty Fellowship #N00014-18-1-2002, and a grant from the Simons foundation collaboration on the global brain. G.H. was supported by The Swartz Foundation.

